# Doblin: Inferring dominant clonal lineages from DNA barcoding time-series

**DOI:** 10.1101/2024.09.08.611892

**Authors:** David Gagné-Leroux, Melis Gencel, Adrian W.R. Serohijos

## Abstract

**Motivation:** The lineage dynamics and history of cells in a population reflect the interplay of evolutionary forces they experience, namely mutation, drift, and selection. When the population is polyclonal, lineage dynamics also manifest the extent of clonal competition among co-existing mutational variants. If the population exists in a community of other species, the lineage dynamics could also reflect the population’s ecological interaction with the rest of the community. Recent advances in high-resolution lineage tracking via DNA barcoding, coupled with next-generation sequencing of bacteria, yeast, and mammalian cells, allow for precise quantification of clonal dynamics in these organisms.

**Results:** In this work, we introduce *Doblin*, an R suite for identifying *do*minant *b*arcode *lin*eages based on high-resolution lineage tracking data. We first benchmarked *Doblin’s* accuracy using lineage data from evolutionary simulations, showing that it recovers clones’ identity and relative fitness in the simulation. Subsequently, we applied *Doblin* to analyze clonal dynamics in laboratory evolutions of *E. coli* populations undergoing antibiotic treatment and in colonization experiments of the gut microbial community. *Doblin’s* versatility allows it to be applied to lineage time-series data across different experimental setups.

**Availability and implementation:** The source code and data are available at https://github.com/dagagf/doblin.

## 1 Introduction

Microbial populations in natural environments or the laboratory often exhibit polyclonality, characterized by genetic differences among individual cells (Lynch and Conery, 2003). This polyclonality is due to their high population size and high rates of new genomic variation, including mutations, recombination, or horizontal gene transfer (Ackermann, 2015; Eyler, 2013). This genomic heterogeneity contributes to phenotypic diversity, such as variable levels of antibiotic tolerance (Balaban, et al., 2004; Lewis, 2007; Shan, et al., 2017), nutrient uptake (Nikolic, et al., 2017; Vilhena, et al., 2018), or growth rates (Kiviet, et al., 2014; Thomas, et al., 2018). Within populations comprised of a single species, polyclonality leads to clonal interference, whereby multiple co-existing clones with beneficial mutations, compete for dominance. This competition may result in the interference or inhibition of the spread of one clone by another, thereby influencing the population’s evolutionary trajectory and dynamics (Desai, et al., 2013). In multiple-species populations, diversity within each species can lead to heterogeneous ecological (inter-species) interactions, with one sub-population of the species competing but another sub-population cooperating with other species in the community (Carrara, et al., 2015; Ferreiro, et al., 2018; Hromada, et al., 2021; Stump, et al., 2022; Wang, et al., 2021). Altogether, accurately quantifying the heterogeneity of communities is crucial for understanding evolutionary and ecological dynamics across diverse scientific domains, including microbial evolution (Good, et al., 2017; Levy, et al., 2015), microbial engineering (Lopez-Garcia, et al., 2010; Rompolas, et al., 2016), and cancer treatment (Landau, et al., 2013; Mroz, et al., 2015).

Recent advancements in chromosomal DNA barcoding technology (Blundell and Levy, 2014), where a unique DNA barcode introduced into the chromosome is transmitted from parent to daughter cells, have enabled high-resolution measurement of clonal lineages in various organisms, including yeast (Blundell and Levy, 2014; Blundell, et al., 2019; Cvijovic, et al., 2018), bacteria (Abel, et al., 2015; Gencel, et al., 2022; Jasinska, et al., 2020; Theodosiou, et al., 2023; Vasquez, et al., 2021), and mammalian cells (Rogers, et al., 2018). In bacteria, these studies led to quantitative analysis of the emergence of antimicrobial resistance, while in human cell lines, they facilitated the quantification of cancer recurrence and metastasis (Ben-David, et al., 2018; Bhang, et al., 2015; Hata, et al., 2016; Nguyen, et al., 2015; Shaffer, et al., 2018). High-resolution lineage tracking of microbial populations also revealed the coexistence of sub-populations exhibiting unique functional and phenotypic characteristics. Diverse fitness levels and growth rates has been observed, potentially influencing cell fate and differentiation into specific lineages (Biddy, et al., 2018; Hollmann, et al., 2020; Lu, et al., 2011; Pei, et al., 2017). Furthermore, lineage tracking has shown that evolutionary events, such as the acquisition of beneficial mutations by a sub-population, can lead to an increase in its frequency and significantly impact the collective behavior of the population and its response to environmental changes (Barrick, et al., 2009; Blundell, et al., 2019; Jasinska, et al., 2020; Levy, et al., 2015; Venkataram, et al., 2016).

A crucial aspect of analyzing the high-resolution dynamics resulting from DNA barcoding is determining the clonal lineages or sub-populations with similar lineage behaviors. However, currently, no computational tools are available to identify dominant barcode lineages from high-resolution lineage tracking data. Existing tools leverage single nucleotide variant data from wholegenome sequencing (WGS) to track and reconstruct sub-population clonal dynamics within tumors (Fischer, et al., 2014; Rubanova, et al., 2022) and not immediately applicable to DNA barcoding data. While tools like *PyFitSeq* (Li, et al., 2018) estimate fitness values of DNA barcodes, but they do not necessarily identify clonal lineages.

To address this gap, we developed *Doblin*, an R suite for quantifying dominant clonal lineages from DNA barcoding time-series data. This package performs clustering of barcode lineage time-series, leveraging the idea that similarities in lineage trajectories reflect comparable relative fitness values. Our approach also identifies persistent clonal lineages in the population. As demonstration, we first applied *Doblin* on data of lineage dynamics from forward evolution simulations, showing its ability to identify lineages with distinct fitness levels. Then, we applied *Doblin* to high-resolution lineage tracking data from a laboratory evolution of *E. coli* under antibiotic resistance and from *E. coli* invasion of the gut microbiome. *Doblin* is an open-source R package available at https://github.com/dagagf/doblin.

## 2 Methods

### 2.1 An overview of the package

*Doblin’s* workflow is shown in Fig. 1. It takes as input high-resolution lineage tracking data (Fig. 1A) obtained from DNA bar-codes of a microbial population. In principle, any time-series data from pooled competition experiments of variants, such as mutational libraries from deep mutational scans (Fowler, et al., 2010; Gray, et al., 2018) or CRISPR screens (Katoh, et al., 2017; Ronda, et al., 2016; Zalatan, et al., 2015), could also be used as input. The input data must be formatted into a table containing barcode identifiers (IDs), timestamps, and read counts associated with each barcode. This data setup is particularly relevant for studies involving serial passaging techniques, where the evolution and propagation of microbes are observed over successive generations.

**Figure 1.**
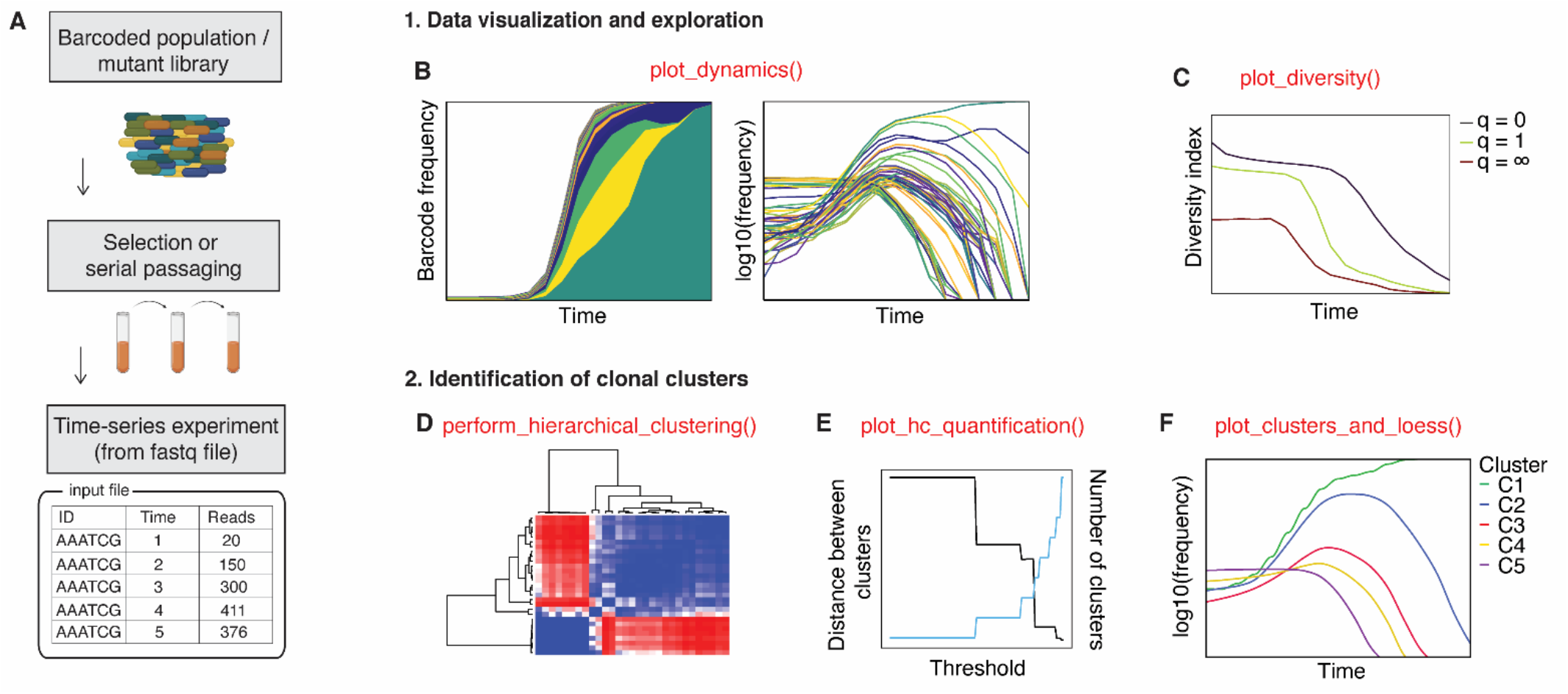
Overview of Doblin. (**A**) Doblin determines the dominant clonal lineages from high-resolution barcoded population time-series data. The input data can also be time-series of mutants from selection experiments of mutational libraries or variants from CRISPR/Cas screens. The flowchart shows the main functions (bold text in red) and steps (bold underlined text). (**B-C**) Tools for data visualization and overview. The function *plot_dynamics()* generates linear and logarithmic representations of the barcode lineage frequencies (panel **B**). The *plot_diversity()* function quantifies the diversity of chromosomal barcodes using the effective diversity index, 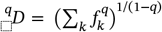 (panel **C**). Diversity calculation is based on the count of unique barcodes (*q* = 0), frequency-weighted barcodes (*q* = 1), and the number of dominant barcodes (*q* = ∞). (**D-E-F**) Tools for identification of dominant clonal lineages: The function *perform_hierarchical_clustering()* compares pairs of frequency trajectories to create a distance matrix based on either Pearson’s correlation or Dynamic Time Warping (DTW). The resulting matrix is used to hierarchically group frequency trajectories with similar behaviors (panel **D**). The function *plot_hc_quantification()* evaluates the clusters produced by hierarchical clustering across a range of thresholds (panel **E**). Once the optimal threshold has been selected by the user, the function *plot_clusters_and_loess()* computes the LOESS average for each cluster at that hierarchical level. Together, the LOESS averages, or “clonal clusters”, reflect the predominant patterns within the dataset (panel **F**). Clonal clusters are ranked according to their final abundance levels.

### 2.2 Data visualization and calculation of effective diversity

*Doblin*’s functionality enables data exploration, whereby the dataset dynamics are visualized with the function *plot_dynamics()*. This function allows users to choose between logarithmic and/or linear scales for plotting barcode frequencies (Fig. 1B). The logarithmic scale emphasizes low-frequency yet persistent cells, while the linear scale highlights dominant clones that have proliferated in the population.

To facilitate the comparison of dominant barcode lineages across biological samples, *Doblin* employs a consistent color mapping strategy: each dominant barcode lineage is assigned a unique color. To implement this strategy, users need to establish a frequency threshold that distinguishes dominant lineages from nondominant ones. The frequency *f*_*i*_(*t*) of lineage *i* at time *t* is calculated as *f*_*i*_(*t*) = *x*_*i*_(*t*)/∑_*k*_ *x*_*k*_(*t*), where *x*_*i*_(*t*) represents the read count of lineage *i* at time *t* within a set of *k* lineages. *Doblin*’s consistent color mapping scheme is intended to ensure clarity and facilitate the interpretation of lineage dynamics under different biological conditions.

Additionally, since barcode diversity reflects the evolutionary and/or ecological pressures encountered by the barcoded population, *Doblin* provides functionality for plotting various diversity metrics using the function *plot_diversity()* (Fig. 1C). This function calculates the effective diversity index 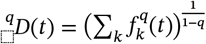 (Jost, 2006; Tuomisto, 2010), where 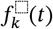 represents the frequency of each barcoded lineage k, and the parameter *q* denotes the diversity index. When *q* = 0, the diversity 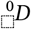 reflects the total number of unique barcodes (or IDs), regardless of their frequencies. This metric corresponds to the “species richness” in ecology, although in the context of barcode dynamics, “species” refers to barcode IDs. When *q* = 1, the diversity index 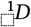 represents the exponential of Shannon diversity. Finally, when *q* = ∞, the diversity index 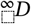 equals the reciprocal of the frequency of the most dominant barcode. By comparing the diversity index across these three orders of *q*, we can effectively characterize the composition of a dataset.

### 2.3 Identification of dominant clonal lineages

#### 2.3.1 Pairwise distance matrix between lineage time series

The second step in *Doblin* is identifying dominant clonal lineages. These clonal lineages are estimated by clustering barcode time series according to their shape and temporal similarities. To improve the accuracy of the clustering, barcode IDs with insufficient time-points are filtered out. This filtering requires users to specify both the number of timepoints to be included and a frequency threshold, ensuring that only dominant and persistent lineages are retained for the downstream analysis.

Identifying the dominant clonal lineages begins with assessing the similarities between pairs of frequency trajectories. This step involves computing a distance matrix. Two methods are available for computing the matrix: one relies on Pearson’s correlation, while the other utilizes Dynamic Time Warping (DTW). Users can choose their preferred method based on experimental data and intended analysis. The primary objective of computing the distance matrix is to hierarchically group frequency trajectories exhibiting similar behaviors. With Pearson’s correlation method, the distance Δ*F*_*xy*_between two frequency trajectories *f*_*x*_ and *f*_*y*_ is computed as:

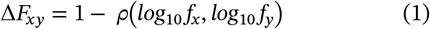

where *ρ*(*log*_10_ *f*_*x*_, *log*_10_ *f*_*y*_) represents the Pearson’s correlation coefficient between the trajectories. A distance close to 0 signifies a positive correlation between the lineages, while a distance nearing 2 indicates negative correlation.

Conversely, DTW adopts a different approach for time-series comparison, prioritizing shape, and overall patterns over point-to-point similarities. It quantifies the alignment between two time series by determining the optimal warping path, thereby accommodating local temporal shifts and variations. The computation of DTW involves creating a cumulative distance matrix *D*(*i, j*) between the i^th^ element of *f*_*x*_ and j^th^ element of *f*_*y*_ defined as:

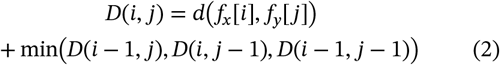

Here, *d*(*f*_*x*_[*i*], *f*_*y*_[*j*]) represents the local distance between the i^th^ element of *f*_*x*_ and j^th^ element of *f*_*y*_, which can be either the Euclidean or Manhattan distance. DTW dynamically computes the cumulative distance matrix and identifies the optimal alignment across all warping paths between the two time-series.

The choice between Pearson’s correlation and DTW depends on the specific characteristics of the time-series and the objectives of the analysis. Pearson’s correlation is preferred when quantifying the strength and direction of the linear relationship between trajectories. On the other hand, DTW is particularly well-suited for capturing nuanced relationships in data exhibiting temporal shifts and non-linear patterns, such as stretching or compression. These patterns are commonly observed in the context of varying growth rates, transients gut dynamics or ecological interactions.

#### 2.3.2 Hierarchical clustering

The calculated distance matrix is used to hierarchically group frequency trajectories exhibiting similar behaviors, as quantified by Pearson’s correlation or DTW. This hierarchical clustering process is carried out by the function *perform_hierarchical_clustering()*, which employs clustering algorithms such as Unweighted Pair Group Method with Arithmetic Mean (UPGMA) or Unweighted Pair Group Method with Centroid Averaging (UPGMC), as provided by the function *stats::hclust()* in R. These clusters encapsulate distinct trajectories, and their characterization is achieved by computing a consensus trajectory via local regression (LOESS). The generated LOESS averages, referred to as “clonal clusters”, represent the dominant behaviors of the dataset and are ranked according to their frequency at the last timepoint (presumably the end of the experiment).

As hierarchical clustering yields a tree structure, determining the number of resulting clonal clusters relies on a cutoff threshold. Traditional clustering evaluation metrics such as the Silhouette Coefficient and Dunn’s Index are often used to determine the optimal number of clusters (Raihan, 2023). However, with time series data, these metrics may not adequately assess clustering quality (Dunn, 2008). In Doblin, we determine the optimal number of clusters by comparing the distance between cluster centroids and the resulting cluster counts across various cutoff thresholds. This is done by the function *plot_hc_quantification()* and is guided by several user-defined considerations. First, we improve robustness against sequencing errors by allowing users to filter out clusters containing fewer than a user-specified number of lineages. Clusters of high-frequency barcodes, even if limited in number of member lineages, are retained. Typically, this is the case when only a few barcodes sweep the population, as shown in the Example #1 below. Secondly, we calculate the distances between clonal clusters by computing the Euclidean distance between their LOESS averages. Our method focuses on identifying the crossover point where the minimum distance between cluster centroids (their LOESS averages) and the overall cluster count converges. Setting the threshold too low may produce many clusters that are similar. Conversely, a high threshold may yield too few clusters, grouping distinct clonal lineage dynamics together. Our approach enables the user to explore the data and select an optimal cutoff threshold.

Below, we demonstrate *Doblin*’s applicability in quantifying dominant clonal lineages across three distinct datasets with increasing level of complexity. First, we apply *Doblin* to evolutionary simulations of a single-species population, where the fitness of each cell is known (Application #1). We show that the clusters identified by *Doblin* indeed correspond to barcodes with similar fitness values. Second, we apply *Doblin* to data from an *in vitro* experimental evolution with high-resolution DNA barcoding of *E. coli* under antibiotic treatment (Application #2). Third, we apply *Doblin* to high-resolution DNA barcoding data of *E. coli* colonizing the mouse gut microbiome with other resident bacterial species (Application #3).

## 3 Results

### 3.1 Application I: Using *Doblin* to analyze high-resolution lineage data from evolutionary simulations

A barcode’s frequency trajectory reflects its fitness level relative to the population’s mean fitness. Barcodes with similar frequency trajectories typically exhibit similar fitness levels. Given that our approach focuses on clustering lineages according to the shape of their trajectories, the clonal clusters generated by *Doblin* are expected to demonstrate varying fitness levels. We benchmarked Doblin’s accuracy using simulation data with known fitness values to validate its effectiveness in extracting dominant clones exhibiting distinct fitness scores. We simulated the evolution of a population of N= 10^7^ barcoded cells (with 10^5^ unique barcodes) using Wright-Fisher process (Gauthier, et al., 2019). Figure 2A illustrates the resulting frequency trajectories throughout the simulation. Initially, all barcoded cells were assigned identical fitness scores (*f* = 1), and each cell’s frequency was tracked over 1125 generations. During the simulation, cells were subjected to *de novo* mutations at a rate of 10^−5^ mutation per genome per replication. The fitness effects of these mutations followed a normal distribution with a mean *μ* = −0.02 and a standard deviation of *σ* = 0.02, indicating a bias towards deleterious mutations (85% deleterious, 15% beneficial). Supplementary Figure 5A shows the frequency trajectories in a Muller plot, depicting their variation over time on a linear scale, thus highlighting the most dominant lineages.

**Figure. 2.**
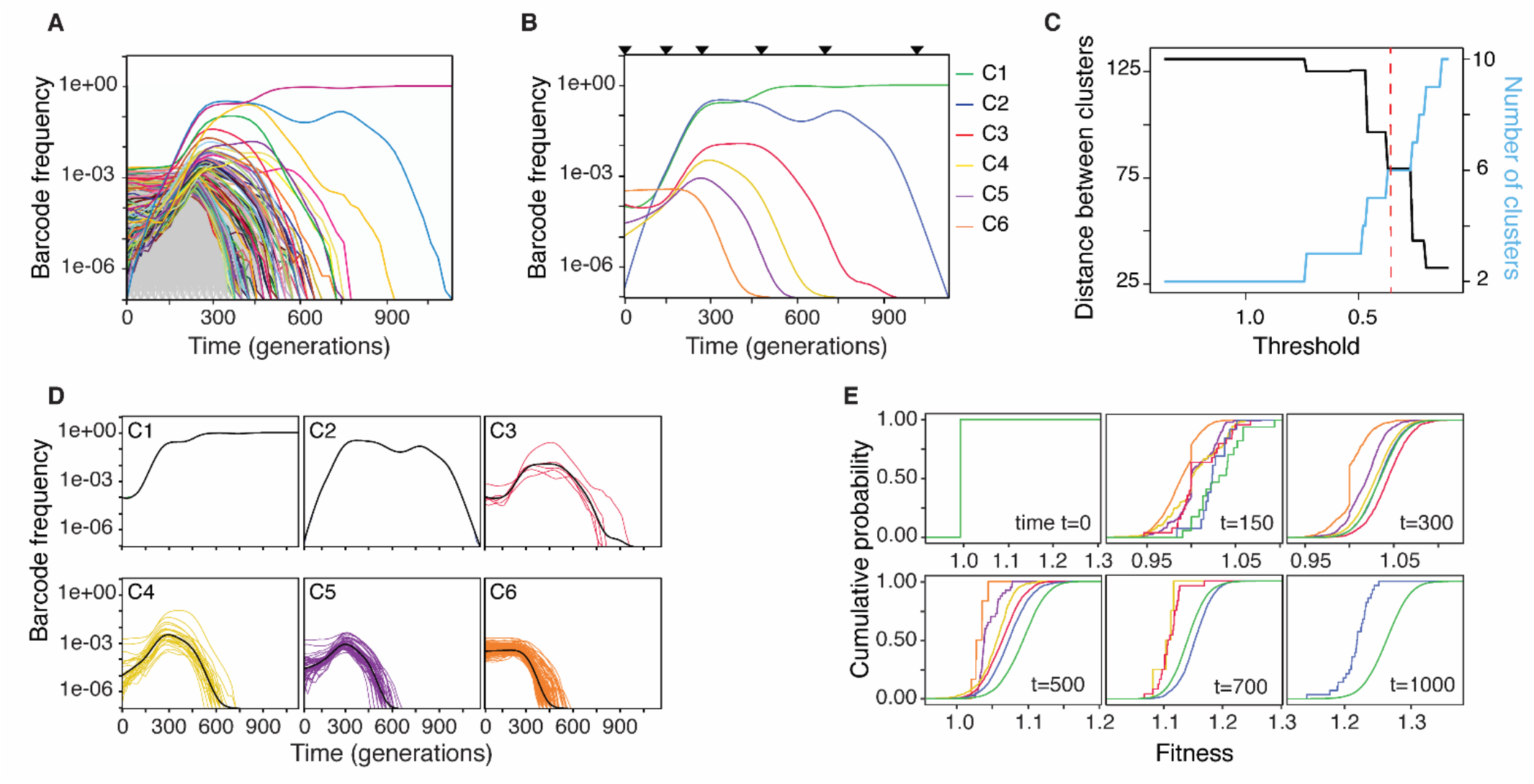
Application #1: Using *Doblin* to analyse high-resolution lineage data from evolutionary simulations. (**A**) Forward evolutionary simulations of bacterial populations using the Wright-Fisher process implemented in SodaPop (Gauthier, et al., 2019). Approximately 10^7^ barcoded cells were initially assigned with equal fitness (*f* = 1). During the evolution, cells acquired *de novo* mutations at a rate of 10^−5^ per genome per replication. The selection coefficient *s* of the mutant is drawn from a normal distribution (mean = -0.02, s.d. = 0.02) and updates the fitness of the mutant as *f*_*mutant*_ = *f*_*mutant*_(1 + *s*). The top 1000 barcodes with a mean frequency over time greater than 10^−4^ are colored uniquely, whereas the rest are shown in gray. (**B**) Consensus trajectories, or centroids, of the 6 clonal clusters identified by *Doblin*. These trajectories correspond to the dominant behaviors of the forward evolutionary simulations. The distribution of fitness across the population for specific timepoints (arrows) are shown in panel E. (**C**) The quantification of hierarchical clustering shows how the Euclidean distance between consensus trajectories (black curve) varies with the number of subsequent clusters (blue curve). The intersection between these two curves (red dashed line) represents the heuristically optimal clustering cutoff, which generated 6 clonal clusters (i.e., C1 – C6). (**D**) Composition of each clonal cluster. These clusters were obtained using a pairwise distance matrix based on Pearson’s correlation. Lineages with mean frequency over time greater than 10^−4^ and persisting for at least 12 time points out of 46 are included in the clustering. The consensus trajectory (black curve) for a cluster is obtained by local regression (LOESS). Note, Cluster 1 (C1) contains the barcode lineage that swept the population (see panel A). (**E**) Cumulative distribution functions (CDFs) of the fitness of all cells in each clonal cluster at the indicated time point. All cells (and clones) have *f* = 1 at the start of the simulation (all CDFs are

To identify the dominant behaviors within the simulated abundance time series, we clustered the frequency trajectories using Pearson’s correlation method. Only the trajectories with an average frequency greater than 10^−4^, and persistent for at least 12 of the 46 timepoints, were retained for clustering. Shown in Figure 2D are the resulting clusters obtained by hierarchical clustering. By choosing a threshold at the intersection of the two criteria (i.e. the cluster count, and the inter-cluster distance) (Fig. 2C), we obtained six distinct groupings (C1 – C6). Examining the composition of each cluster showed that lineages with similar shapes were grouped together (Fig. 2D). Cluster 1 (C1 – green trajectory) contained the most frequent barcode that swept the population. Cluster 2 (C2 – blue trajectory), the second most frequent clonal cluster, initially mirrored C1’s trajectory before diverging around the 450th generation—C1 acquired a beneficial mutation that allowed it to outcompete C2. Clusters C3 to C6, despite beginning with frequencies on par with C1, experienced extinction at different intervals throughout the simulation. At different timepoints in the simulation (arrows in Fig. 2B), we show the distribution of fitness for cells in each clonal cluster (Fig. 2E). At time *t* = 0, all clones had identical fitness levels. However, as time progressed, each clonal trajectory diverged due to their differences in fitness. Clonal clusters with lower fitness levels began to go extinct by the 400th generation. While clonal clusters C1 and C2 demonstrated similar fitness distributions and frequency trajectories until the 450th generation, a beneficial mutation within Cluster C1 increased its fitness within the population. This mutation eventually reached fixation, allowing C1 to dominate the population by the end of the simulation. In contrast, Cluster C2 maintained a constant fitness level before experiencing a decrease, followed by a temporary increase at the 750th generation due to another mutation. However, this mutation did not confer a significant advantage over C1, resulting in the eventual extinction of C2. Altogether, these results demonstrate that *Doblin*’s inferred clonal clusters are distinguishable by their fitness scores, as expected by a shape-based clustering approach.

We also applied Doblin to evolutionary simulations with different levels of poly-clonality and clonal interference by varying the extent of standing genetic variation and effects of *de novo* mutations (Supplementary Figures 1 to 3). Overall, the clonal clusters identified by *Doblin* capture lineages with distinct fitness in these diverse evolutionary scenarios.

### 3.2 Application II: Using *Doblin* to extract dominant lineage behaviors from an experimental evolution of *E. coli*

We applied *Doblin* to high-resolution lineage data from *in vitro* evolution of *E. coli* under the antibiotic Trimethoprim (TMP) (0.1 µg ml^-1^ TMP, replicate 2) with a population size of ∼3×10^7^ cells (Jasinska, et al., 2020). Unlike simulated data, the frequency trajectories derived from experimental data exhibit more complex evolutionary dynamics (Figure 3A, Supplementary Figure 5B). Since *Dolin*’s aim is to estimate dominant lineage behaviors, we restricted our analysis to include only the lineages with a mean frequency exceeding 10^−4^ and persisting for at least 12 consecutive time points out of the 16. These cut-offs can be modified by the user to explore the effects of experimental noise that could strongly affect the dynamics of low-frequency barcodes. In Figure 3B, we show the composition of the resulting clonal cluster, indeed revealing clusters that group barcodes with similar dynamics. The clonal lineage dynamics (Fig. 3A) is indicative of the presence of clonal interference. Indeed, certain lineages initially benefiting from positive selection (i.e. C4, C5 and C6) diminish later as higher-fitness competitors emerge (i.e. C1 and C2). Furthermore, the initial increase observed in C1, C4 and C6 may be caused by pre-existing beneficial mutations present at the start the experiment (standing genetic variation). After 100 generations, while the frequencies of C4 and C6 began to decline, C1 emerged as the dominant cluster. This observation suggests that mutations within C1 underwent selection, providing a competitive advantage to lineages within this cluster. In contrast, the frequencies of C2 and C3 increased later in the experiment, accompanied by significant fluctuations. This pattern indicates that lineages within these clonal clusters likely acquired *de novo* mutations which, unlike those in C1, failed to pass selection.

**Figure 3.**
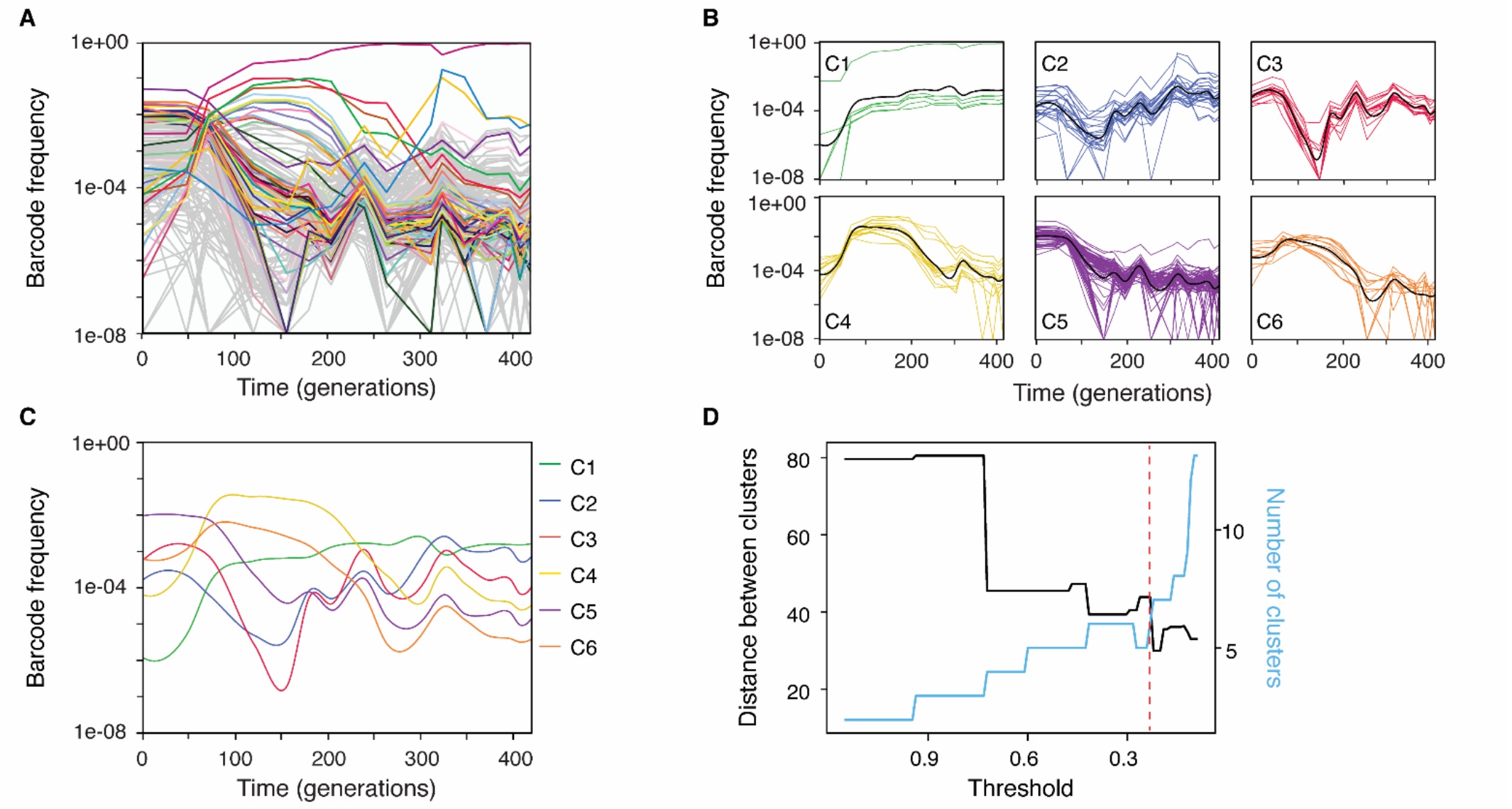
Application #2: *Doblin* applied to a month-long experimental evolution of *E. coli* under antibiotics. (**A**) Frequency trajectories of barcoded lineages throughout a month-long evolution of E. coli under antibiotic (Trimethoprim = 0.1 µg ml^-1^, replicate 2 in (Jasinska, et al., 2020)). The top 50 barcodes with a mean frequency over time greater than 10^−4^ are uniquely colored, whereas the rest are shown in gray. (**B**) Composition of the clonal clusters identified by *Doblin*. These clusters were obtained using a pairwise distance matrix based on Pearson’s correlation. Only the lineages with mean frequency over time greater than 10^−4^ and persisting for at least 12 out of 16 time points were retained. Colored lines correspond to unique chromosomal barcodes in the cluster, while each cluster’s black line denotes its LOESS average. The clonal clusters are ordered based on their average frequency at the final time point. Here, cluster 1 (C1) contains the barcode that manifested the clonal sweep. (**C**) LOESS averages, or consensus trajectories, of the 6 clonal clusters identified by *Doblin*. They reflect the dominant behaviors observed in the dataset. (**D**) The threshold (red dashed line) indicates the heuristically optimal clustering, resulting in 6 clusters (i.e. C1 – C6).

We also applied Doblin to compare the dominant clonal dynamics in two antibiotic concentrations, one at TMP = 0.1 µg ml^-1^ and another that has 1 order of magnitude lower (TMP = 0.01 µg ml^-1^) from (Jasinska, et al., 2020). While genetic drift contributed to the loss of several individual low-frequency lineages in both conditions, it commenced later in the lower TMP condition (Supplementary Figure 4). The rate of barcode diversity loss is expected to correlate with the strength of selection pressure, which is the antibiotic concentration in this case. Consequently, the lower TMP concentration led to slower diversity collapse, leading to a longer period of time where clonal interference occurs. Supplementary Figure 4 demonstrates these expected behaviors, where the clonal cluster dynamics exhibit a slower sweep and more complex dynamics in the lower TMP concentration.

### 3.3 Application III: Using *Doblin* to extract dominant behaviors from abundance time series of *E. coli* invading the gut microbiome of mouse

Laslty, we applied *Doblin* to high-resolution lineage data obtained from a two-week-long *E. coli* evolution experiment in mice gut to demonstrate the effect of increased population heterogeneity on population dynamics (Gencel, et al., 2022). A notable observation is the clones that arrive first in the gut are not always the guaranteed “winner” (dominant clone) at the end of the experiment (Supplementary Figure 5C). By examining the clonal clusters, we can observe that the first barcodes to enter the gut are either removed from population or segregate at a lower frequency (Figure 4B and Figure 4C). This observation highlights the influence of transient dynamics of gut colonization.

**Fig. 4.**
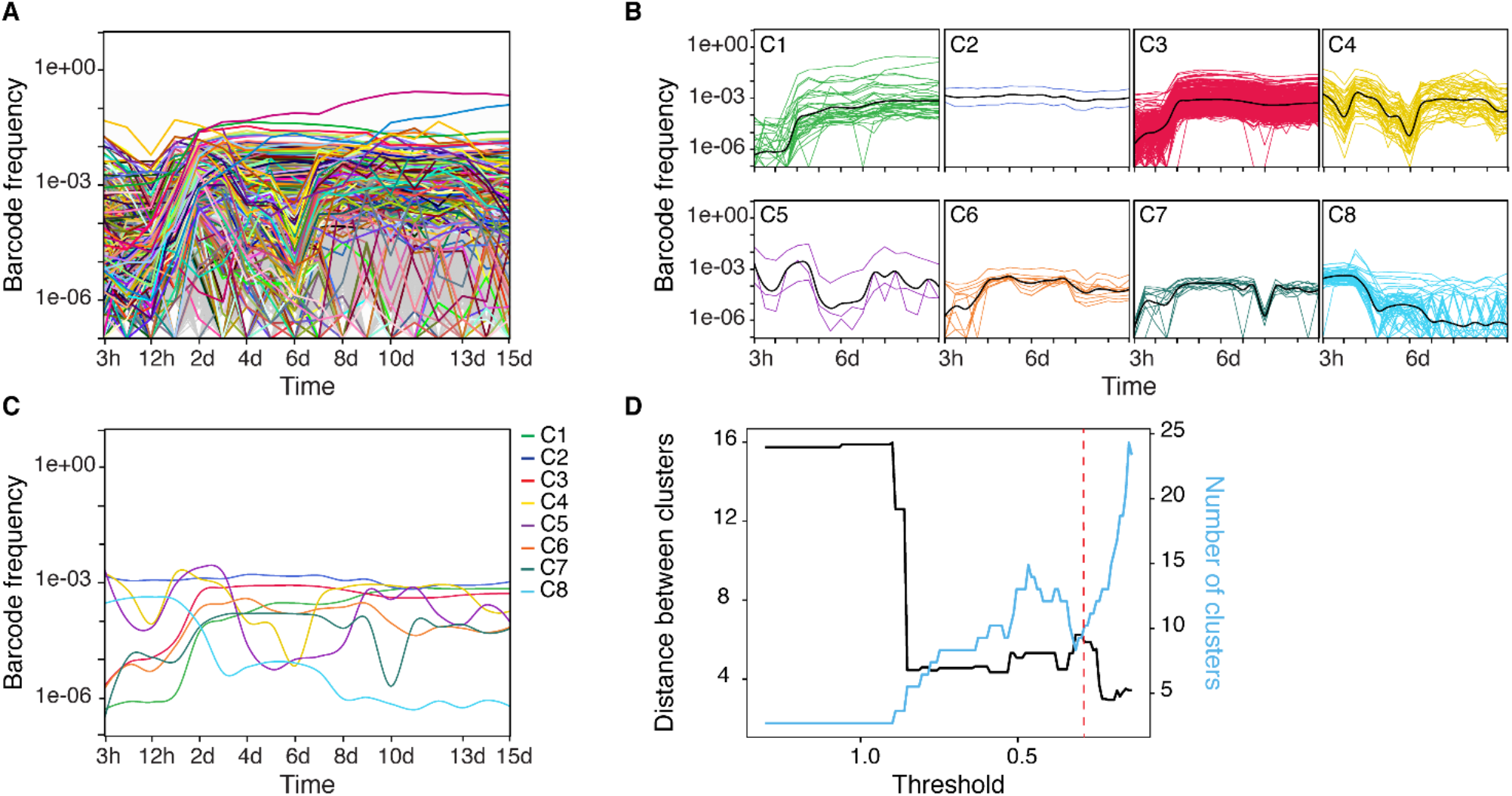
Application #3: Using *Doblin* to extract dominant behaviors from abundance time series of *E. coli* invading a mouse gut microbiome. (**A**) Frequency of ∼10^8^ chromosomal barcoded cells of *E. coli* over 2-week period in mice gut with pre-existing microbiome (from rm1 of cohort 2) (Gencel, et al., 2022). The top 1000 barcodes with a mean frequency over time greater than 5×10^−4^ are colored uniquely, whereas the rest are shown in gray. (**B**) Composition of the clonal clusters identified by *Doblin* using a pairwise distance matrix based on Pearson’s correlation. We retained only the lineages with mean frequency over time greater than 5×10^−5^ and persistent for at least 10 out of 18 time points. (**C**) The LOESS averages of the 8 identified clonal clusters correspond to the dominant behaviors of the dataset. (**D**) The threshold (red dashed line) corresponds to the heuristically

The dominant clonal cluster, C1, demonstrates better adaptation to the gut environment. Clonal clusters C4 to C8 persist at lower frequencies and exhibit significant fluctuations. These variations could reflect ecological contributions, such as interactions with other species within the microbiome, or the emergence of *de novo* mutations. However, by analyzing the clonal cluster information in conjunction with 16S rRNA gene sequencing data, we can quantitatively infer the ecological interaction dynamics (Gencel, et al., 2022). Figure 4 shows that Doblin is able to recover the dominant clonal cluster from inherently complex abundance time-series.

### 3.4 Computational time

Runtime varies with input size and complexity. To assess this metric, we benchmarked key pipeline steps (*filterData, perform_hierarchical_clustering, filterHC, plotHCQuantification*, and *plot_clusters_and_loess*) using experimental and simulated datasets described in this paper on both Windows (Intel i7-1255U, 16GB RAM) and Linux (Intel Xeon E5-2643 v3) systems. For the simulated dataset (306336 barcodes, 17 timepoints, 35 MB), these steps took ∼40 CPU seconds on Windows and were twice as fast on Linux. For the experimental dataset (511280 barcodes, 17 timepoints, 60 MB), they were completed in ∼90 CPU seconds on Windows and ∼40 CPU seconds on Linux.

## 4. Discussion

Here, we introduce *Doblin*, a tool designed for the quantitative analysis of polyclonal communities from high-resolution time series data. *Doblin* distinguishes itself by quantifying the fitness of these communities through their evolutionary temporal dynamics, unlike other available tools that primarily rely on fold enrichment for data analysis. To ensure *Doblin*’s accessibility and applicability for a wide range of users, we have provided detailed documentation of its features and utilities.

*Doblin*’s ability to accurately identify clonal clusters relies on the quality of experimental data. Doblin does not automatically correct for experimental DNA barcode artifacts, which could arise from efficiency in genomic extraction or PCR jackpotting. Additionally, our tool primarily focuses on dominant lineages, potentially overlooking low-frequency barcodes that may become extinct early in the experimental timeline. However, users have the flexibility to adjust parameters to include or exclude certain lineages. The quality of the resulting cluster is also expected to improve with better data, such as higher sampling of the DNA barcode time-series.

## Supplementary information

Supplementary data are available at *Bioinformatics* online.

## Acknowledgements

We thank Shimon Bershtein for discussions and Louis Gauthier for help in the initial development of Doblin. We also acknowledge members of the Serohijos and Bershtein labs for testing the Doblin pipeline.

