## Supporting Figures for "Doblin: Inferring dominant clonal lineages from DNA barcoding time-series"

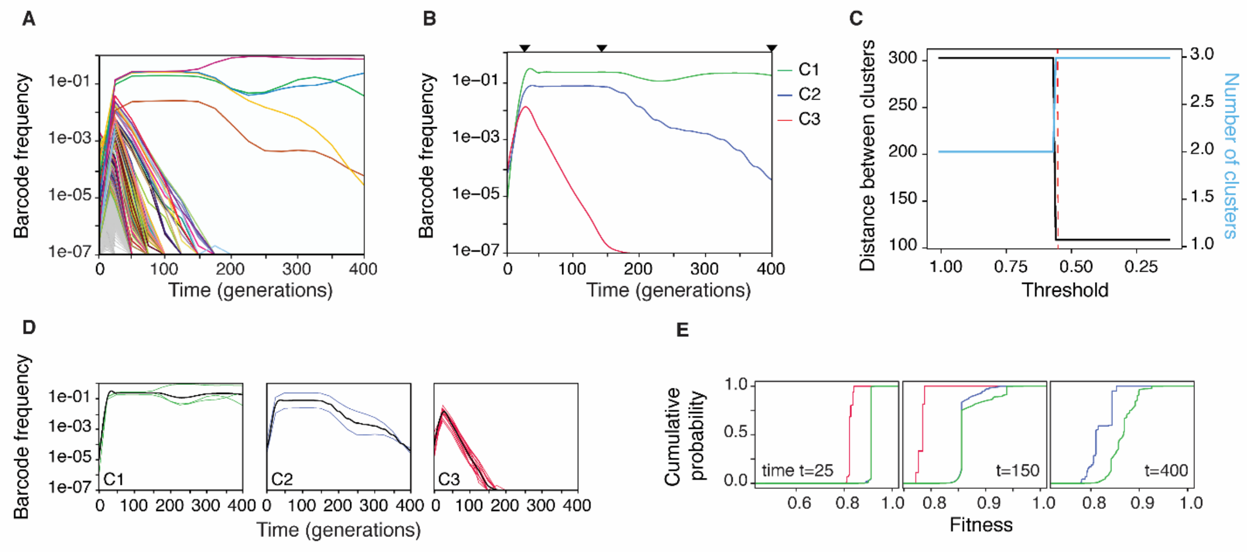


**Supplementary Figure 1. Application of *Doblin* to evolutionary simulations with standing genetic variation and *de novo* mutations.** (**A**) Similar to Figure 2, but cells exhibit different fitness levels at $t = 0$ due to pre-existing mutations, with a mean fitness effect of $s = 0.1$. During the evolution, cells acquired *de novo* mutations at a rate of 2.5×10$^{-4}$ per genome per replication. These new mutations have selection coefficients that follow a Gaussian distribution, with 90% of the values being negative (indicating detrimental effects on fitness) and 10% being positive (indicating beneficial effects). The mean selection coefficient for the beneficial mutations is $s = 0.01$. Lineages with a minimum frequency of 5x10$^{-5}$ are colored uniquely, while all other lineages are in gray. (**B**) The 3 clonal clusters identified by *Doblin*. The distribution of fitness across the population for specific timepoints (arrows) are shown in panel E. (**C**) The quantification of hierarchical clustering shows how the Euclidean distance between clonal clusters (black curve) changes depending on the number of subsequent clusters (blue curve). The intersection between these two curves (red dashed line) represents the heuristically optimal clustering, which generated 3 clonal clusters (i.e., C1 – C3). (**D**) Composition of each clonal cluster. We included lineages in our analysis only if they demonstrated a mean frequency greater than 5×10$^{-5}$ and were present for a minimum of 2 out of 17 time points. (**E**) Cumulative distribution functions (CDFs) of the fitness of all cells in each clonal cluster at the indicated time point. *Doblin* and the clonal clusters group barcoded cell lineages of similar fitness.

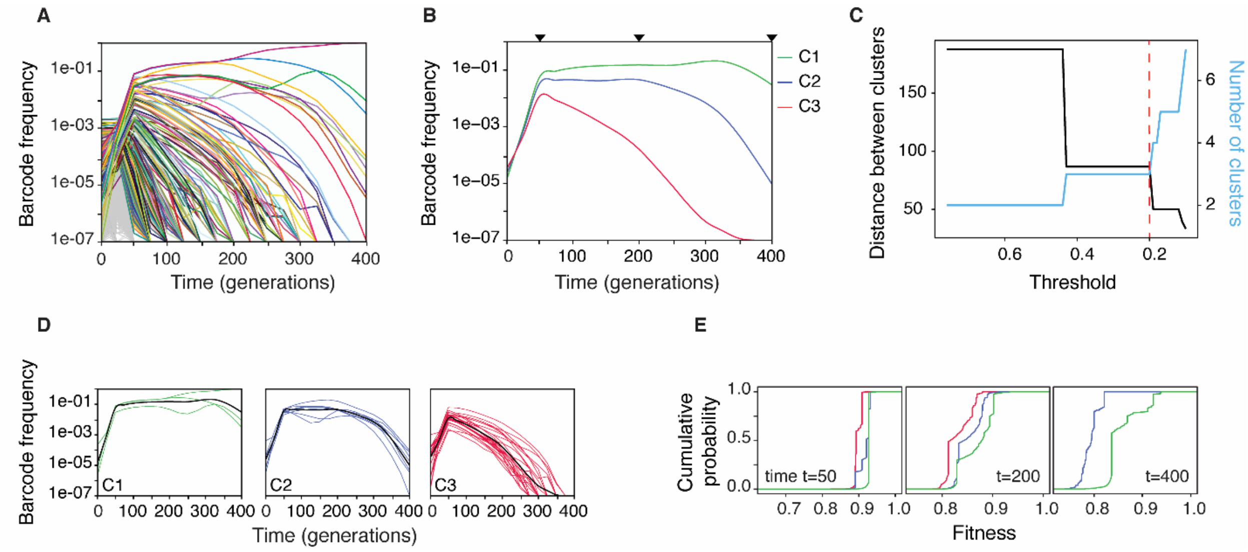


**Supplementary Figure 2. Application of *Doblin* to evolutionary simulations with standing genetic variation and *de novo* mutations.** (**A**) Similar to Supplementary Figure 1, but exhibiting pre-existing mutations with a mean fitness effect of $s = 0.05$. (**B**) The 3 clonal clusters identified by *Doblin*. (**C**) Criteria for finding the optimal number of clusters. (**D**) Composition of each clonal cluster. We included lineages in our analysis only if they demonstrated a mean frequency greater than $5\times10^{-5}$ and were present for a minimum of 10 out of 17 time points. (**E**) Evolution of CDFs of the fitness levels for each clonal cluster. Color descriptions for for B-E are similar to Figure 2 and Supplementary Fig. 1.

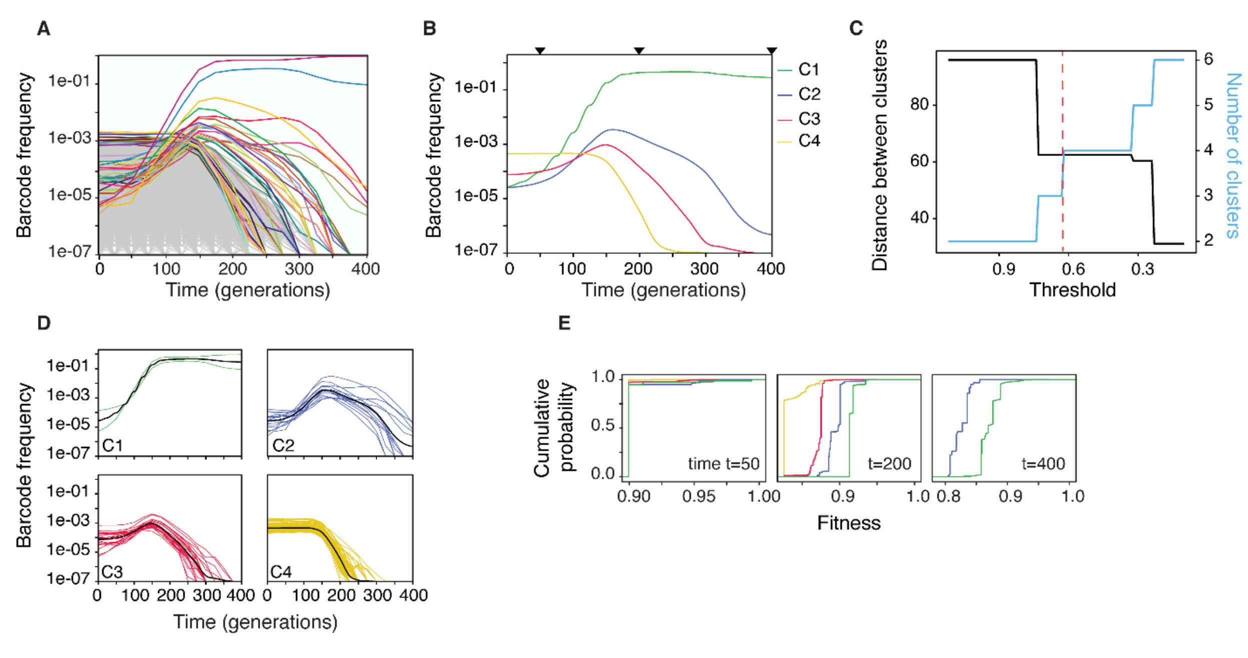


**Supplementary Figure 3. Doblin applied to evolutionary simulations with standing genetic variation and *de novo* mutations.** (**A**) Similar to Supplementary Figure 1, but exhibiting pre-existing mutations with a mean fitness effect of $s = 0.01$. During the evolution, cells acquired *de novo* mutations at a rate of $2.5\times10^{-4}$ per genome per replication. These new mutations have selection coefficients that follow a Gamma distribution, with 100% of the distribution being positive. The mean selection coefficient for the beneficial mutations is $s = 0.01$. Lineages with a minimum frequency of $10^{-4}$ are colored uniquely, while all other lineages are in gray. (**B**) Consensus trajectories of the 4 clonal clusters identified by *Doblin*. (**C**) Criteria for finding the optimal number of clusters. (**D**) Composition of each clonal cluster. We included lineages in our analysis only if they demonstrated a mean frequency greater than $5\times10^{-5}$ and were present for a minimum of 8 out of 17 time points. (**E**) Evolution of CDFs of the fitness levels for each clonal cluster.

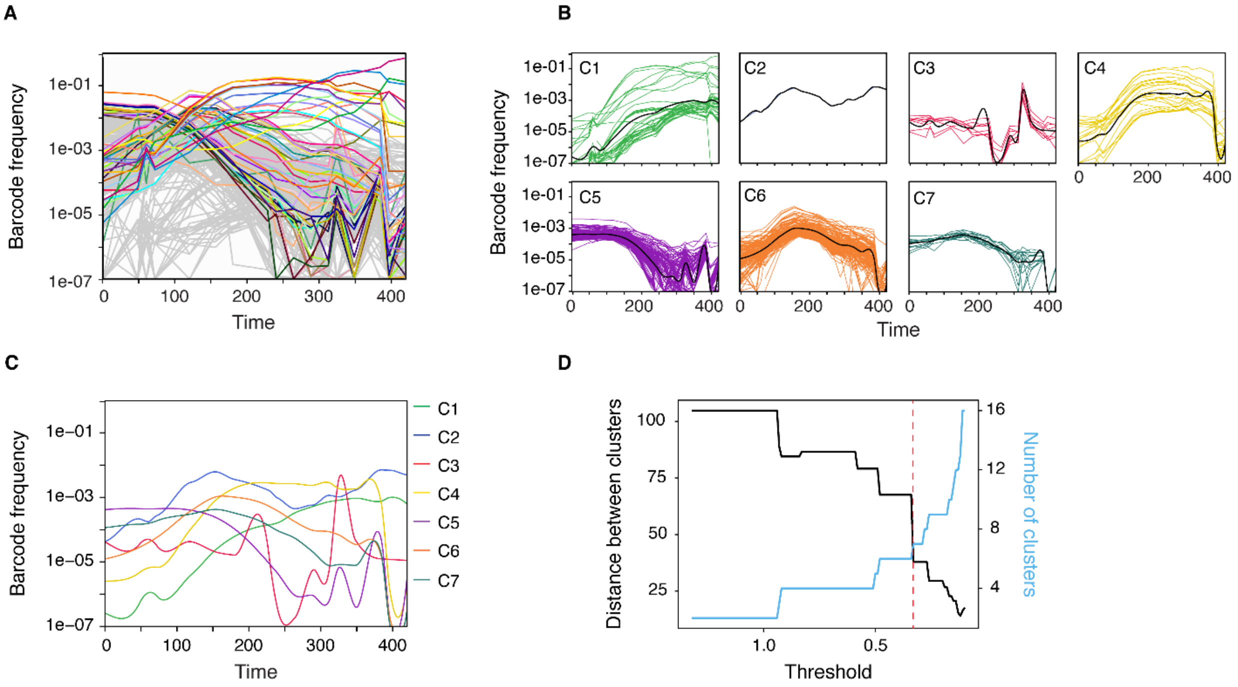


**Supplementary Figure 4. *Doblin* applied to a month-long experimental evolution of E. coli under lower antibiotic regime.** (**A**) Similar to Fig. 3, but under 0.01 μg ml$^{-1}$ of Trimethoprim (replicate 3) (Jasinska, et al., 2020). (**B**) Composition of the clonal clusters identified by *Doblin*. Only the lineages with mean frequency over time greater than 10$^{-4}$ and persisting for at least 12 out of 17 time points were retained for the analysis. (**C**) LOESS averages of the 7 clonal clusters identified by *Doblin*. (**D**) Threshold (red dashed line) for optimal clustering, resulting in 7 clusters (i.e. C1 – C7).

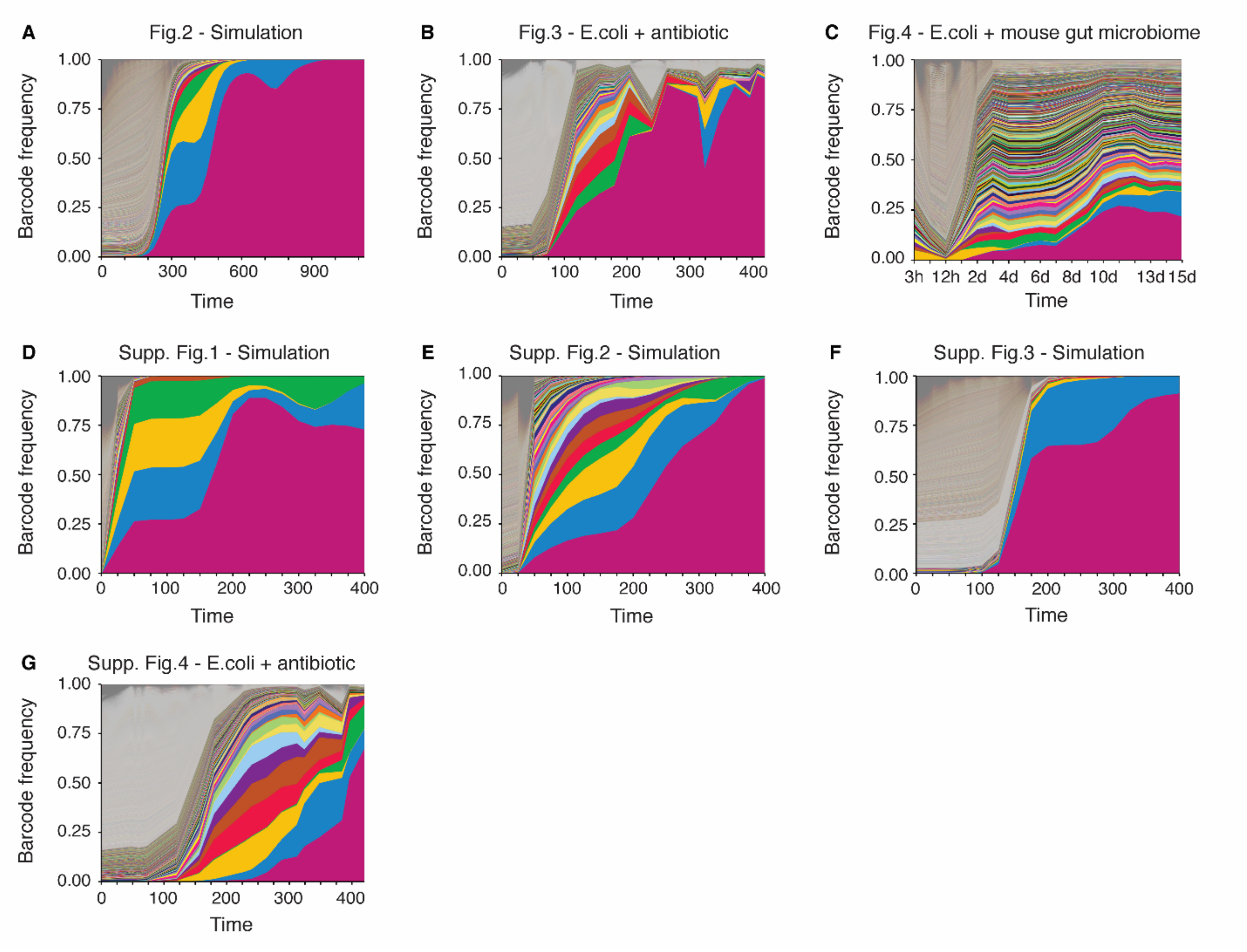


**Supplementary Figure 5. Dynamics of barcoded lineages frequencies in simulation and experiments to which *Doblin* has been applied.** Each panel illustrates frequency trajectories represented as Muller plots, showcasing changes in frequencies over time on a linear scale. The colors used in these plots maintain consistency between the logarithmic and linear scale representations. Panel (**A**) displays the Muller plot of frequency trajectories in Fig. 2A, while Panels (**B**) through (**G**) represent the frequency trajectories of Fig. 3, Fig. 4, and Supplementary Figures 1 through 4, respectively.
